# Effect of telomerization on the growth patterns of adult cutaneous diploid fibroblasts

**DOI:** 10.1101/2021.04.01.437571

**Authors:** Alexander A. Zgurskiy, Leonid Yu. Prokhorov

## Abstract

The article describes effects of telomerization on proliferative activity in cultured cutaneous diploid fibroblasts obtained from an adult men aged 57 years. It is shown that the cells in the control culture initially divide quite actively, but starting from the 27th to the 40th passages, the rate of their division decreases and the time to reach the monolayer increases by about 2 times, and starting from the 42nd passage, the time to reach the monolayer increases by 4-10 times (up to 16-40 days). In a single case, an increase in the doubling time of control non-thelomerized cells was recorded up to 30 times (136 days). The introduction of the telomerase gene allowed cells maintain a high rate of cell division to about 45-47 passages. At the same time, the culture with the telomerase gene exceeds the control culture by 10 passages. Rate of growth the cells with telomerase gene after 48 and up to 57 passages slowed down by 10-18 times (up to 40-73 days). At the 47th passage in the control culture and at the 57th passage in the experimental culture, the cells practically stopped dividing and their number did not increase. The results show that the inclusion of the telomerase gene in adult fibroblasts does not always cancel the limit of cell divisions.

## Introduction

In the 60s of the last century, L. Hayflick and co-authors (Hayflick, 1965; Hayflick and Moorhead, 1961) found that normal somatic cells in culture can divide only a limited number of times. At the same time, it turned out that the number of cell divisions depends on the type of animal from which they are obtained. The number of cell divisions from short-lived species is less than that of cells derived from long-lived species.

Later, the other scientist A. M. Olovnikov (Olovnikov, 1973) from Russia revealed the reason for limiting the number of cell divisions in culture. It turned out that with each division, the end section of the DNA – the telomere is shortened by several nucleotides, and finally, there comes a time when the length of this section is reduced so much that copying the entire DNA chain becomes impossible. As a result, the division of the cell nucleus is not carried out and the cell itself does not divide.

Later it was found that the limit of cell division in culture usually coincides with the average number of divisions that the cells of the body make during its growth, starting from the zygote to its complete formation (Prokhorov, 1999).

If the cell contains the enzyme telomerase, which is based on reverse transcriptase, then it completes the telomeric portion of the DNA to its original size during division and the restriction of cell proliferation does not occur. In normal somatic cells, such as human skin fibroblasts, telomerase is absent. If the telomerase gene is artificially introduced into the DNA of cells, the synthesis of the telomerase enzyme in the cell will begin.

In 1998, A. G. Bodnar et al. (1998) showed for the first time a practical increase in the number of doubling of the culture of human eye epithelial cells by 20 passages above the usual number of divisions after the introduction of the telomerase enzyme gene into their DNA. In 2003, E. E. Egorov et al. (2003) found that the introduction of the telomerase gene into the DNA of adult skin cells significantly expands the proliferative potential and in their experiments they have already achieved 200 doubling of these cells, with 50-60 doubling in the control.

As is known, in culture, somatic human fibroblasts slow down their division at the late passages after about 50 divisions. Hayflick and co-authors conditionally assumed that the culture died when, after re-seeding in a ratio of 1:2, the number of cells in it did not double in 2 weeks. In fact, it was suspected that not all the cells in the culture actually died and the survivors continue to divide, but their rate of division is greatly slowed down. In our experiments, we were interested to learn how intact cell cultures and cultures with cells that have an artificially implanted telomerase gene behave after a conditionally accepted replicative death, that is, when the cells cannot double in 2 weeks.

## Materials and methods

We used human diploid skin fibroblasts (HDSF) obtained from a man (POL) aged 57 years. The cell culture was obtained in the company “Medbiokor” LLC, Russia. To obtain the initial culture, a piece of skin was taken from the forearm in a sterile manner and placed in a Petri dish with a type II collagenase solution. After the cells were released from the biopsy material, a DMEM/F12 1:1 growth medium (PanEco, Russia) with 10% of embryonic calf serum (Biosera, France) was added to the dish. Next, the cells were cultivated in a plastic flasks (SPL, Korea) of 25 cm^2^ on the same medium. After the cells filled the entire surface of the flask, they were re-seeded into a new flask (1:2). Then they were subcultivated after 3-4 days. Thus, 3 subcultivations were performed. Then the cells were subcultivated into borosilicate glass flasks with a external diameter of 54 mm, the area of the internal growth surface is 19.6 cm^2^. The flasks were hermetically sealed with black rubber stoppers, and the cultures were grown in a thermostat at 37°C. On the 7th passage, HDSF cells were telomerized. For this purpose, a 2 ml solution with a lentiviral construct with gene hTERT under CMV promotor was prepared on a growth medium without serum.

The growth medium from the flask with HDSF of 6^th^ passage was poured out, the cells were rinsed with a medium without serum, then 1.5 ml of a medium without serum, 0.5 ml of a solution with a lentiviral structure with gene hTERT and 0.1 ml of a polybren solution were poured to the final concentration of the latter 8 µg/ml. After 1.5 hours, a 4-5 ml full growth medium with serum was poured into the flask and left overnight. The next day, this medium with the lentivirus construct and polybren was poured out, washed with a new growth medium with serum, and then filled with 5-6 ml of the new growth medium with serum. Then cells were subcultivate in the usual way 1:2. We used 3 flasks for the experimental culture and 3 flasks for the control culture. Old flasks were periodically replaced with new flasks during subcultivating. At first, when the cells were rapidly dividing, 1:4 was re-seeded in the early passages, and later, when the rate of cell division slowed down, they began to re-seeded again in the ratio of 1:2 both in the experiment and in the control. The lentiviral construction and polybren were obtained from the Engelhardt Institute of Molecular Biology (Russia). Non − telomerized control cells were designated as HDSF, and telomerized cells were designated as HDSF hTERT.

The cells were counted and photographed using an inverted Biolam P-1 microscope with video output to a computer. The number of cells on the 5 fields of the flask was calculated, the data was averaged, multiplied by the corresponding coefficient, and the number of cells per cm^2^ was obtained.

## Results

The results are shown in Figures 1-12. During 3 months up to passage 27, the control and telomerized cells grew quite rapidly and doubled in 3-4 days, with a maximum of 11-14 days occasionally. From the age of 4 months, they began to notice that normal and telomerized cells began to grow more slowly. But the control cells slowed down more. At 6 months of culture growth, we found a real lag in the number of divisions of non-telomerized cells from telomerized cells by 4-5 passages. At this time, telomerized cells were located at passage 32 and doubled in about 1-1. 5 weeks (7-10 days), while non-telomerized cells were “stuck” at passage 27-28 and doubled in an average of 1-2 weeks. (7-14 days).

**Figure 1.**
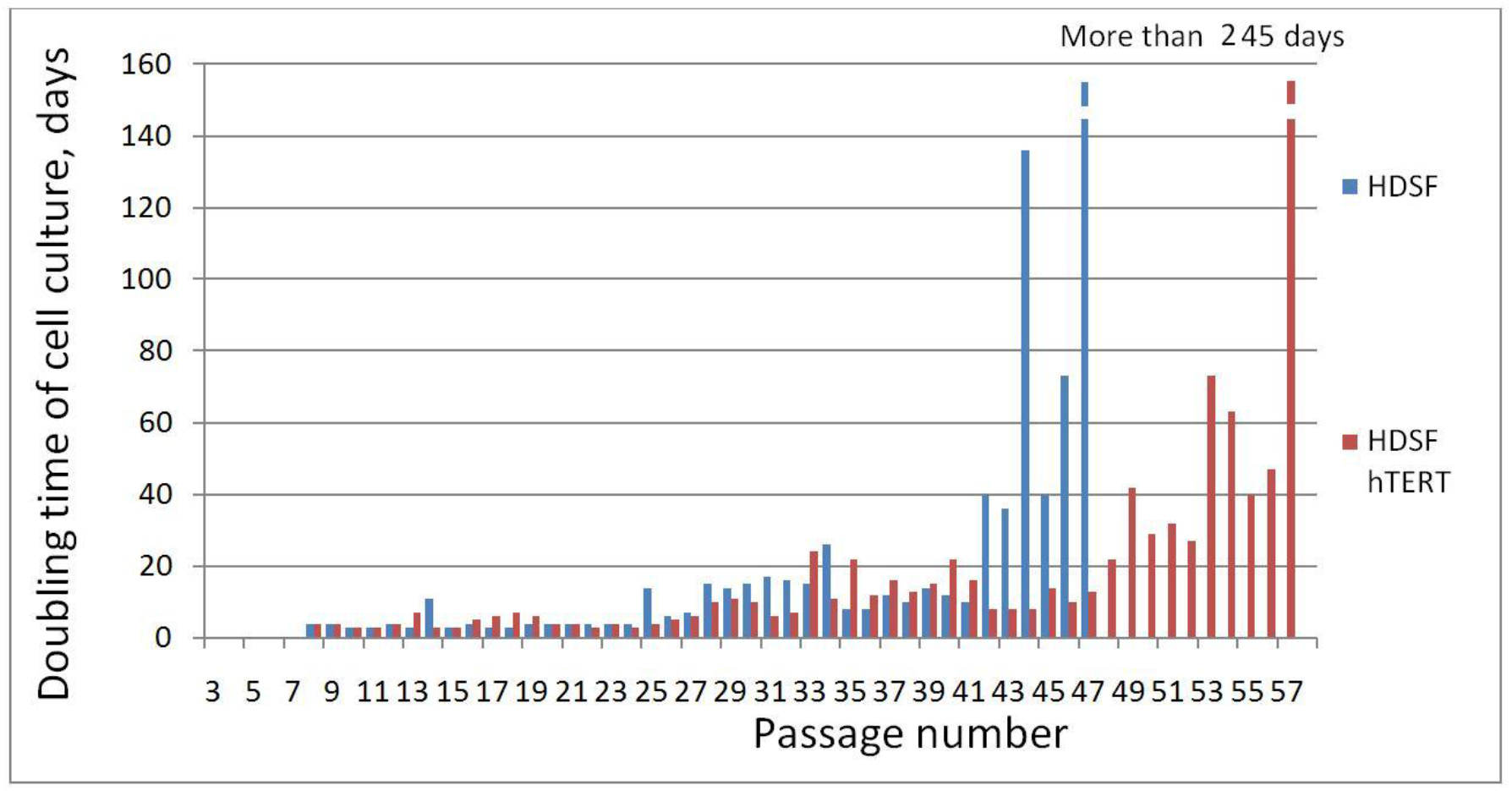
Change in the doubling time of HDSF cell cultures with and without the embedded telomerase gene (HDSF hTERT). Telomerization was performed on the 7th passage.

Since that time, both cultures have slowed down their growth, but the control cells began to divide even more slowly than the telomerized ones and continued to lag behind in the number of passages.

After 15 months, the control cells made 41 passages, and the telomerized cells already made 49 passages, that is, 8 passages more, as can be seen in Figure 1. After 19 months, the gap was already 10 passages (44-47 passages were made by the control culture and 54-57 passages-by the culture with telomesized cells).

Figures 2-3 show photos of cell cultures with and without the telomerase gene on the 10th day of growth after re-seeding, with the control cells located at 41 passage (Figure 2), and the cells with the telomerase gene at a later 47 passage (Figure 3). Despite the larger passage, the culture with telomerized cells has already reached the monolayer, and the control culture is halfway through, which indicates slow cell division, many of which are spread out on the surface and are not ready for division.

**Figure 2.**
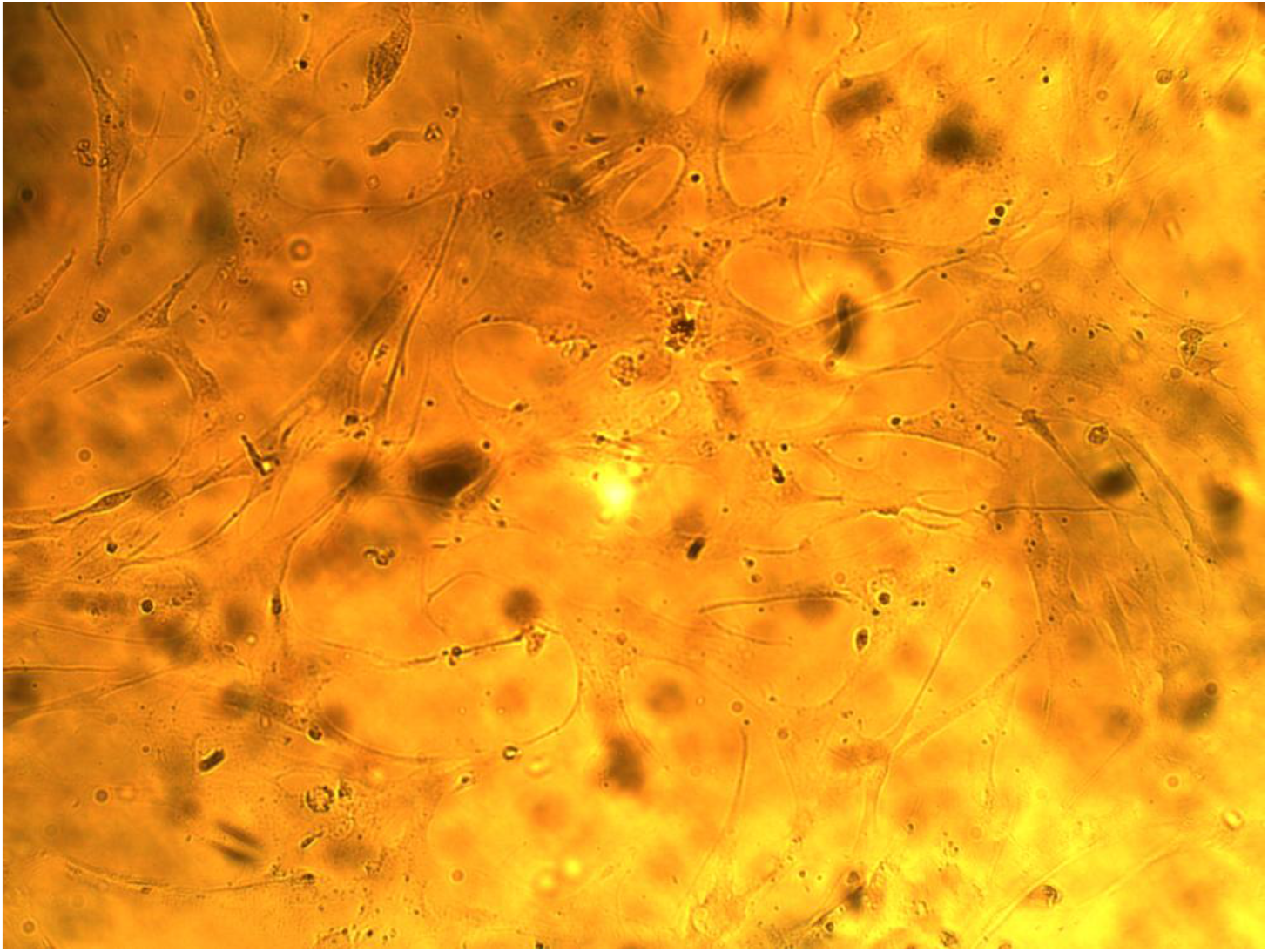
HDSF cells for 41 passage. Growth of 10 days after re-seeding. The density of the culture is 1880 cells / cm^2^.

**Figure 3.**
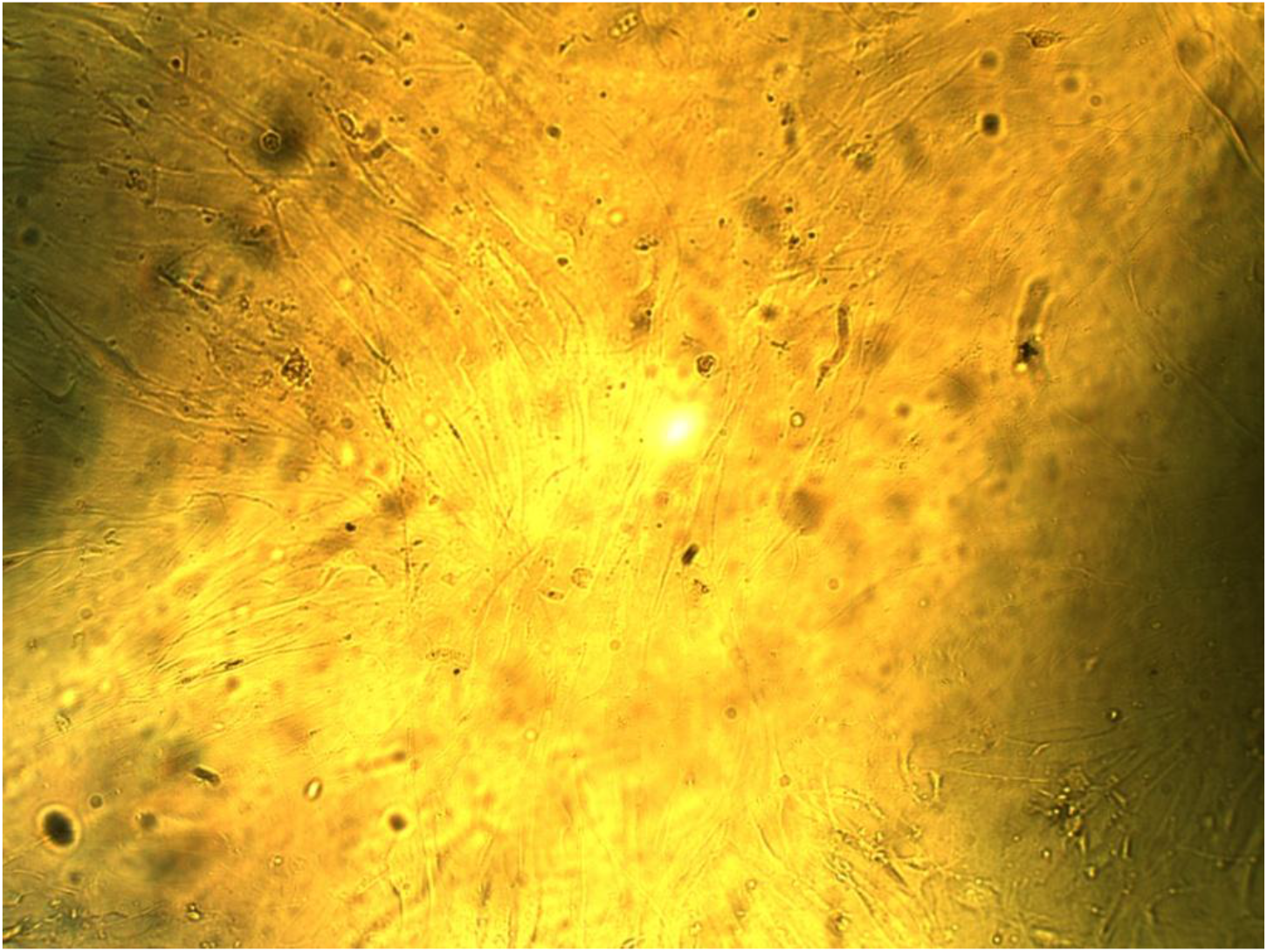
HDSF hTERT for 47 passage. Growth of 10 days after re-seeding. The density of the culture is 24,000 cells / cm^2^.

Figure 4 shows the cells of the HDSF hTERT on the 55th passage after 7 days of growth after re-seeding. The density of the culture is 6450 cells / cm^2^. In Figure 5, the same cells on the same 55 passage after 37 days of growth after re-seeding. The density of the culture is 14800 cells / cm^2^.

**Figure 4.**
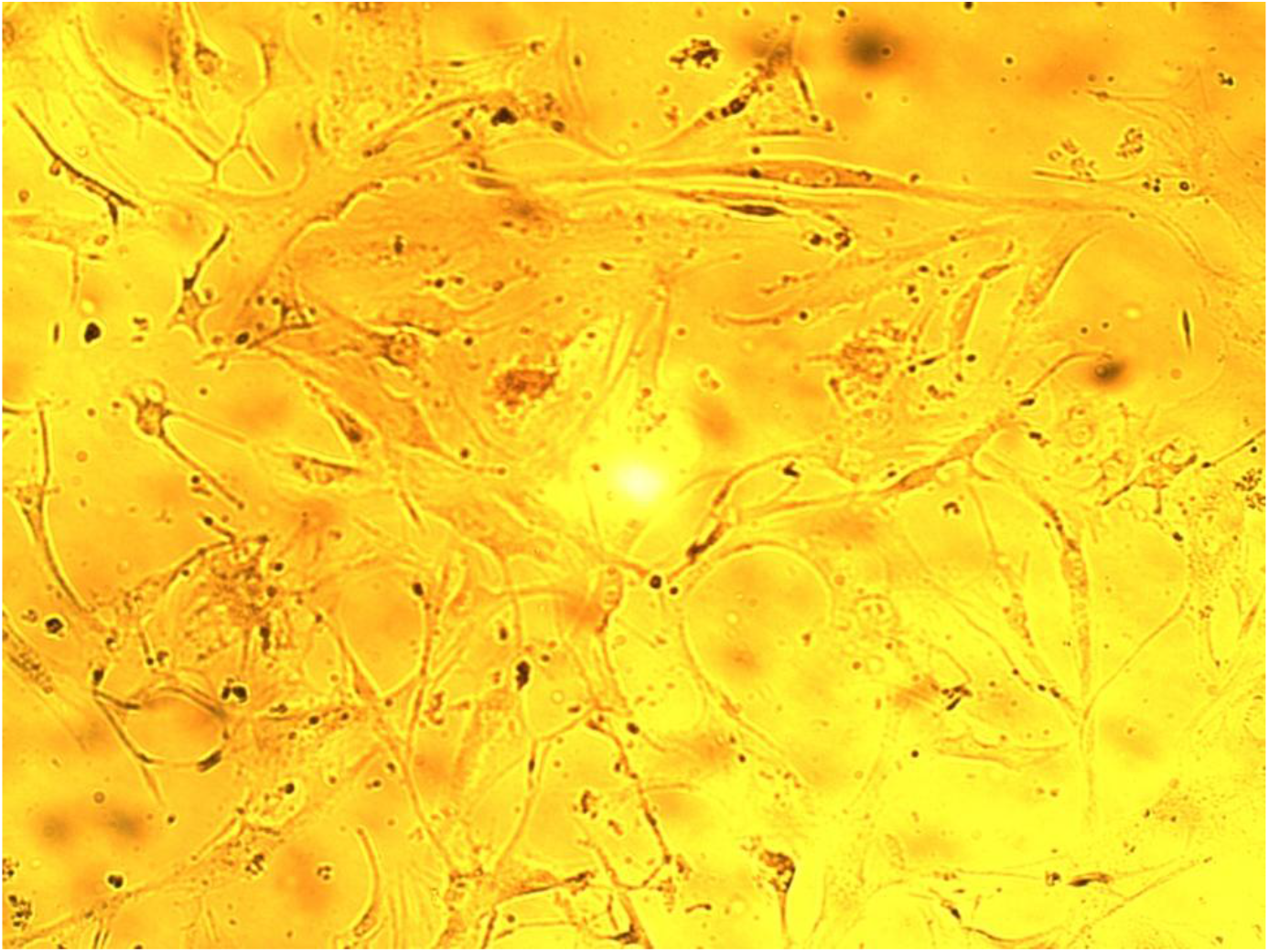
HDSF hTERT for 55 passage. Growth of 7 days after subcultivation. The density of the culture is 6450 cells / cm^2^.

**Figure 5.**
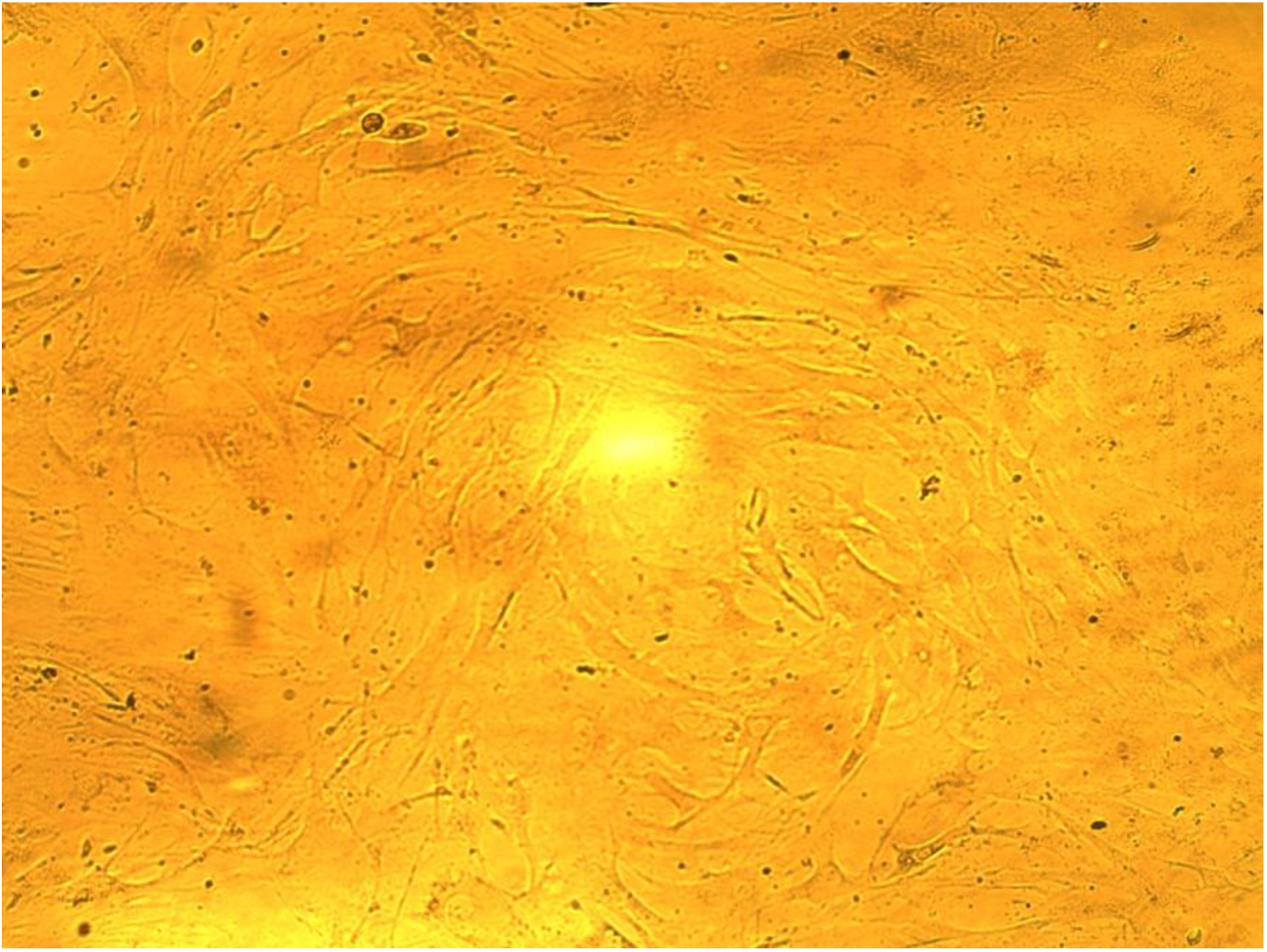
HDSF hTERT for 55 passage. Growth of 37 days after re-seeding. The density of the culture is 14800 cells / cm^2^.

Figure 6a and 6b show non-telomerized HDSF cells on the 45th passage and on the 37th day of growth after re-seeding. Photos of two different places of the same flask are shown. It can be seen that the non-telomerized cells on the 45th passage, which lagged behind the telomerized cells by 10 divisions on the 37th day of growth, have a low density and did not reach the monolayer, the average density is 3800 cells / cm2.

**Figure 6a.**
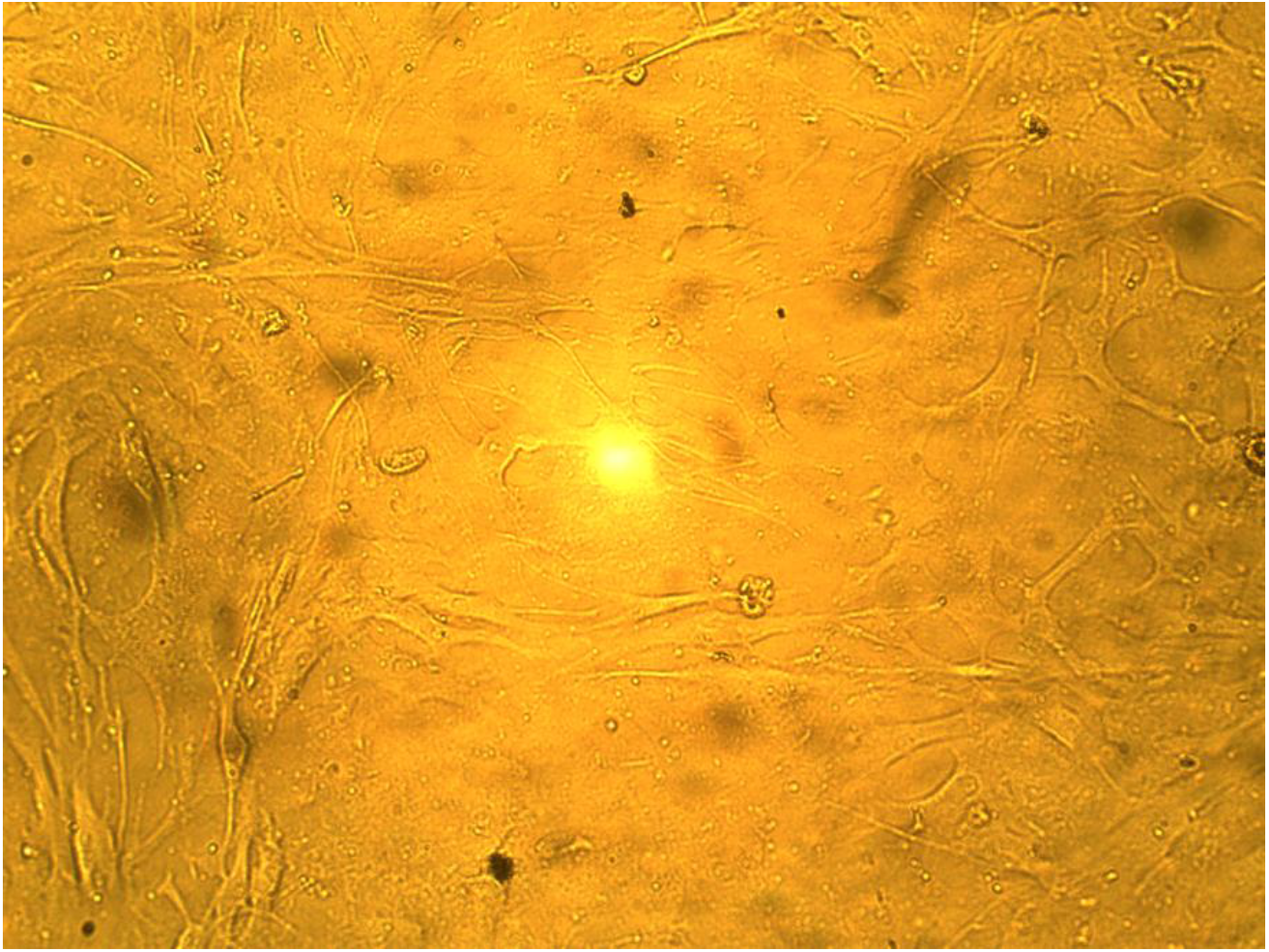
DHSF cells for 45 passage. Growth of 37 days after re-seeding. Avarege density of the culture is 3800 cells / cm^2^. Photo of one of the places in the flask.

**Figure 6b.**
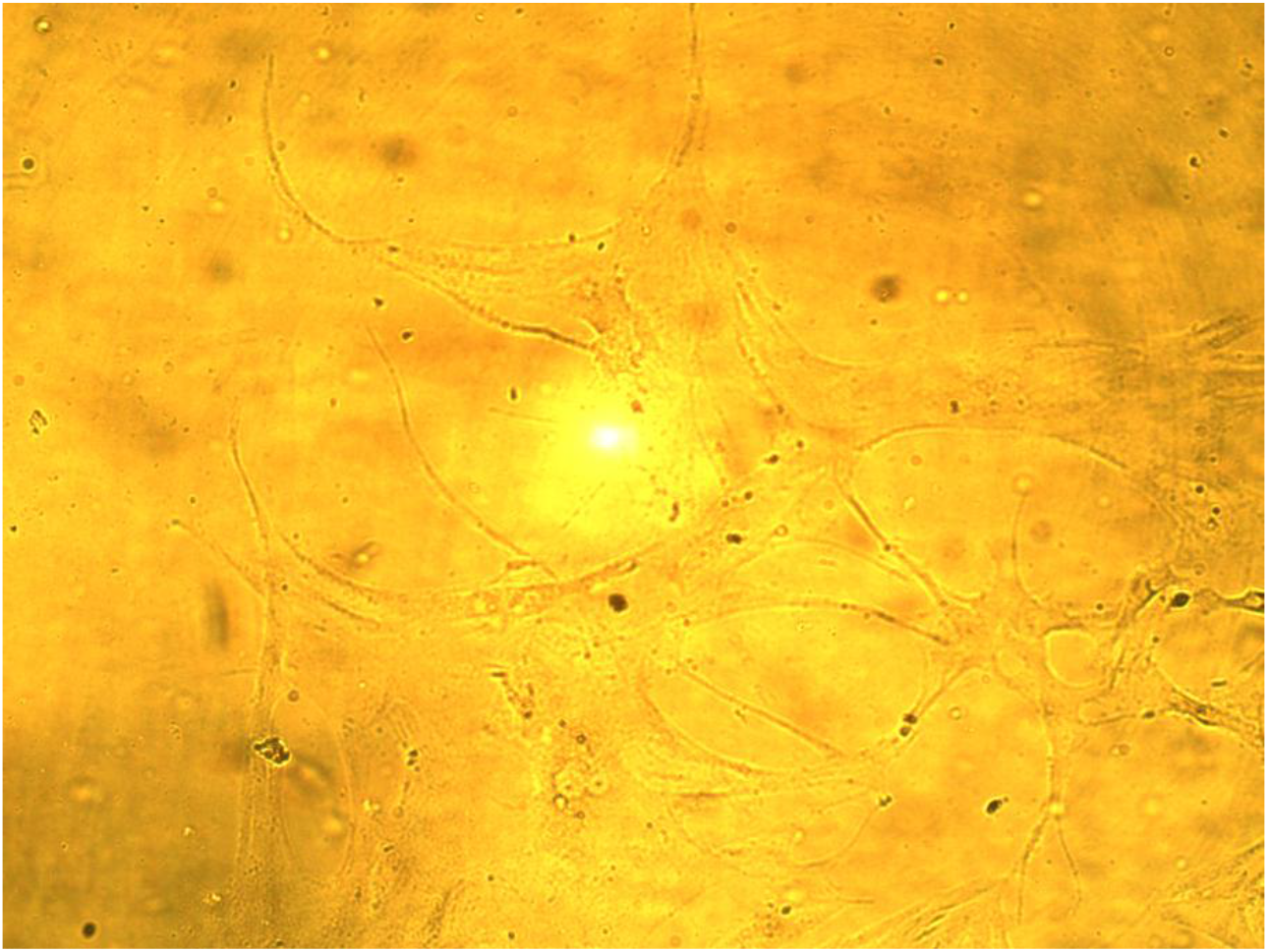
HDSF cells for 45 passage. Growth of 37 days after subcultivation. Avarege density of the culture is 3800 cells / cm^2^. Photo of another place in the same flask.

It is desirable to compare the type of cells in relatively early and late passages. In the early passages (20-30), when the cells are still dividing quite quickly, they have the appearance of a spindle, tightly pressed together. Figure 7 shows the control HDSF on the 23rd passage, the growth of 21 days after re-seeding. The cells are located close to each other, have the appearance of a flow in the same direction and do not intersect with each other.

**Figure 7.**
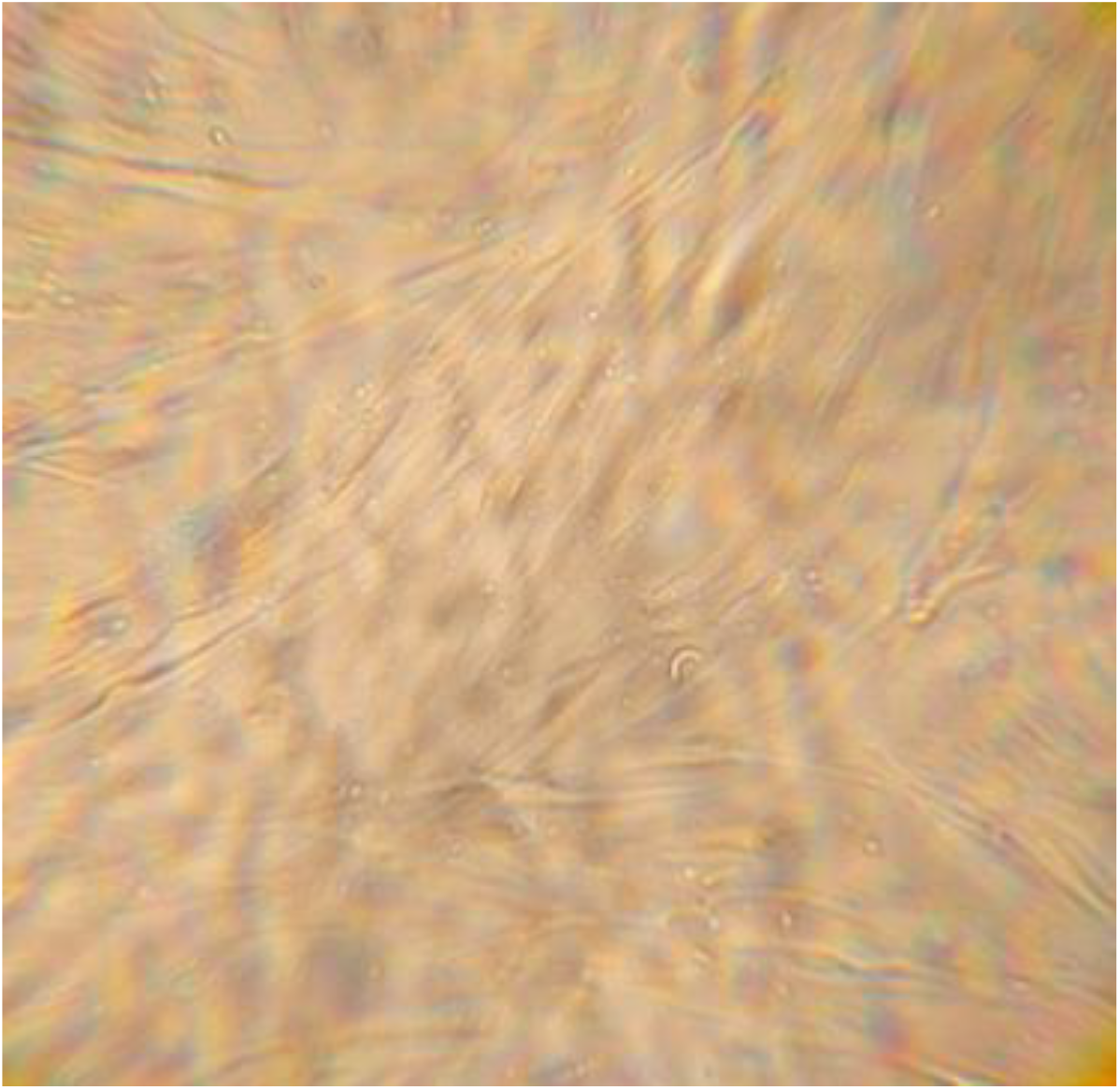
HDSF cells 23 passage, 21 days after replanting. The average density of the culture is 35 000 cells / cm^2^.

The view of the control cells in passage 45, the growth of 52 days after re-seeding is shown in Figures 8a, 8b, and 8c. Fibroblasts are more branched, lie farther apart, have an uneven density in different parts of the flask, and are randomly directed.

**Figure 8a, b. c.**
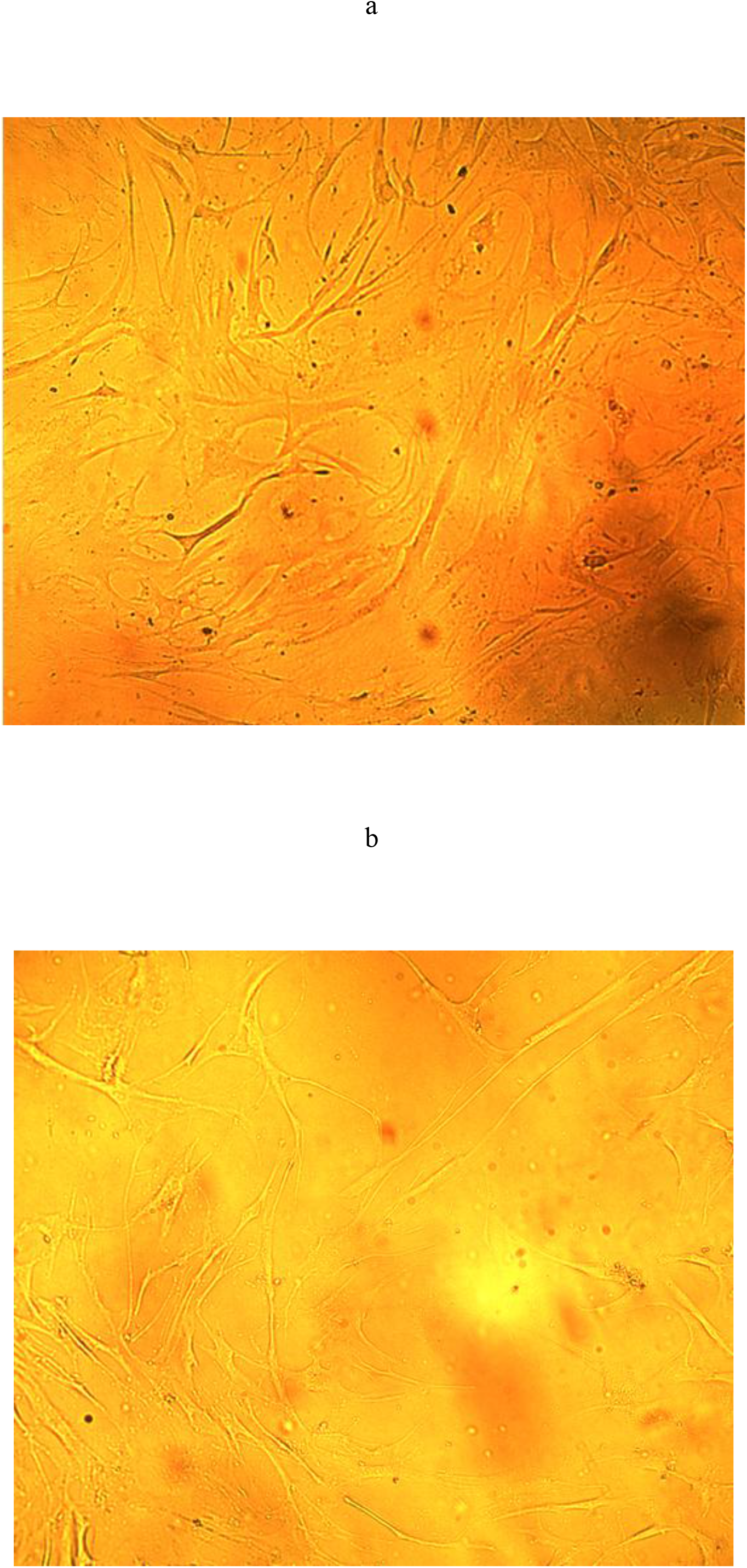
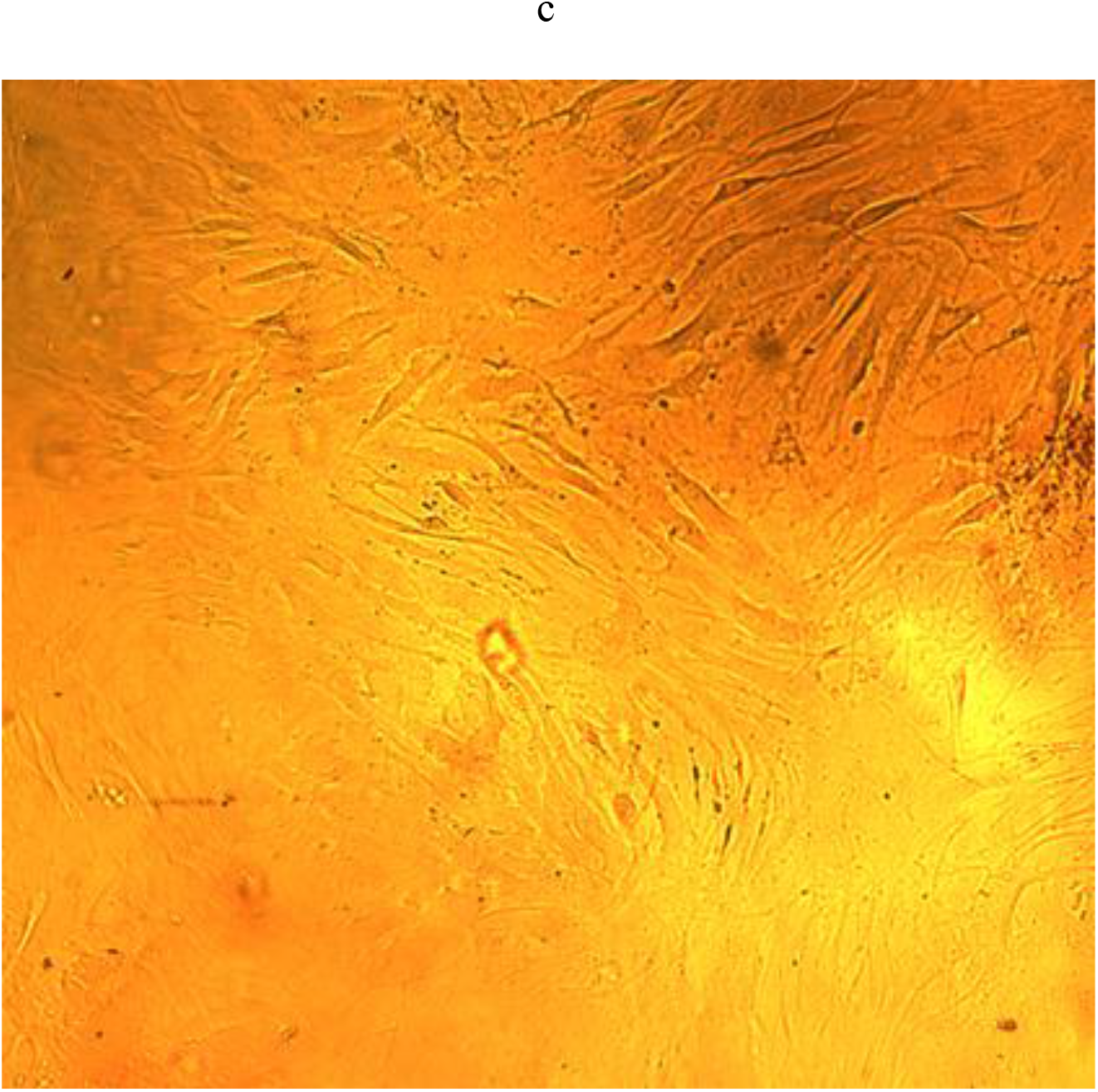
HDSF cells 45 passage, 52 days after re-seeding. Different zones of the same flask. The average density of the culture is 18000 cell / cm^2^.

The telomerized cells on the 29th passage, growth 11 days after re-seeding are shown in Figure 9. As in the control “young” culture, the cells are in good condition, lie side by side, have an elongated shape, and there are no outgrowths.

**Figure 9.**
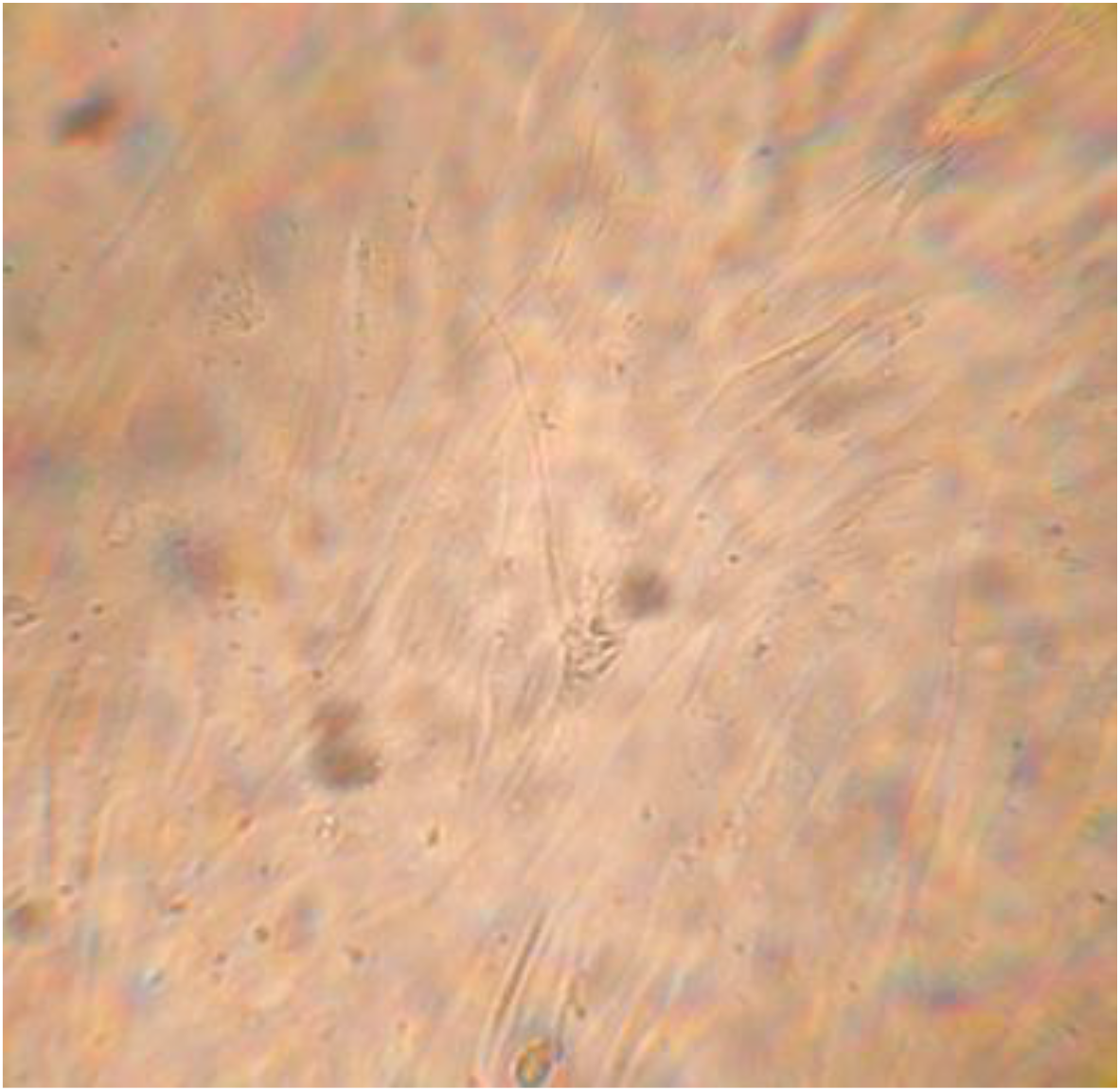
HDSF hTERT 29 passage, growth 11 days after re-seeding. The average density of the culture is 36,000 cells / cm^2^.

The telomerized cells on the 55th passage, growth 40 days after re-seeding, are shown in Figure 10. It can be seen that many cells in the culture are in a state of hypertrophy, their shape is uneven. At this time, the cells not only increase in size in area (horizontally), but also become less voluminous, their height decreases, as evidenced by the blurring of the cell boundaries, they seem to spread out on the growth surface. This is also indicated by the difficulty in pointing the microscope at sharpness, because the cells merge with the background. This is more noticeable in telomerized cells of passage 57 after 33 days of growth after re-seeding (Figure 11). In some fibroblasts, many outgrowths appear in an amount of up to 6-7-10 or more.

**Figure 10.**
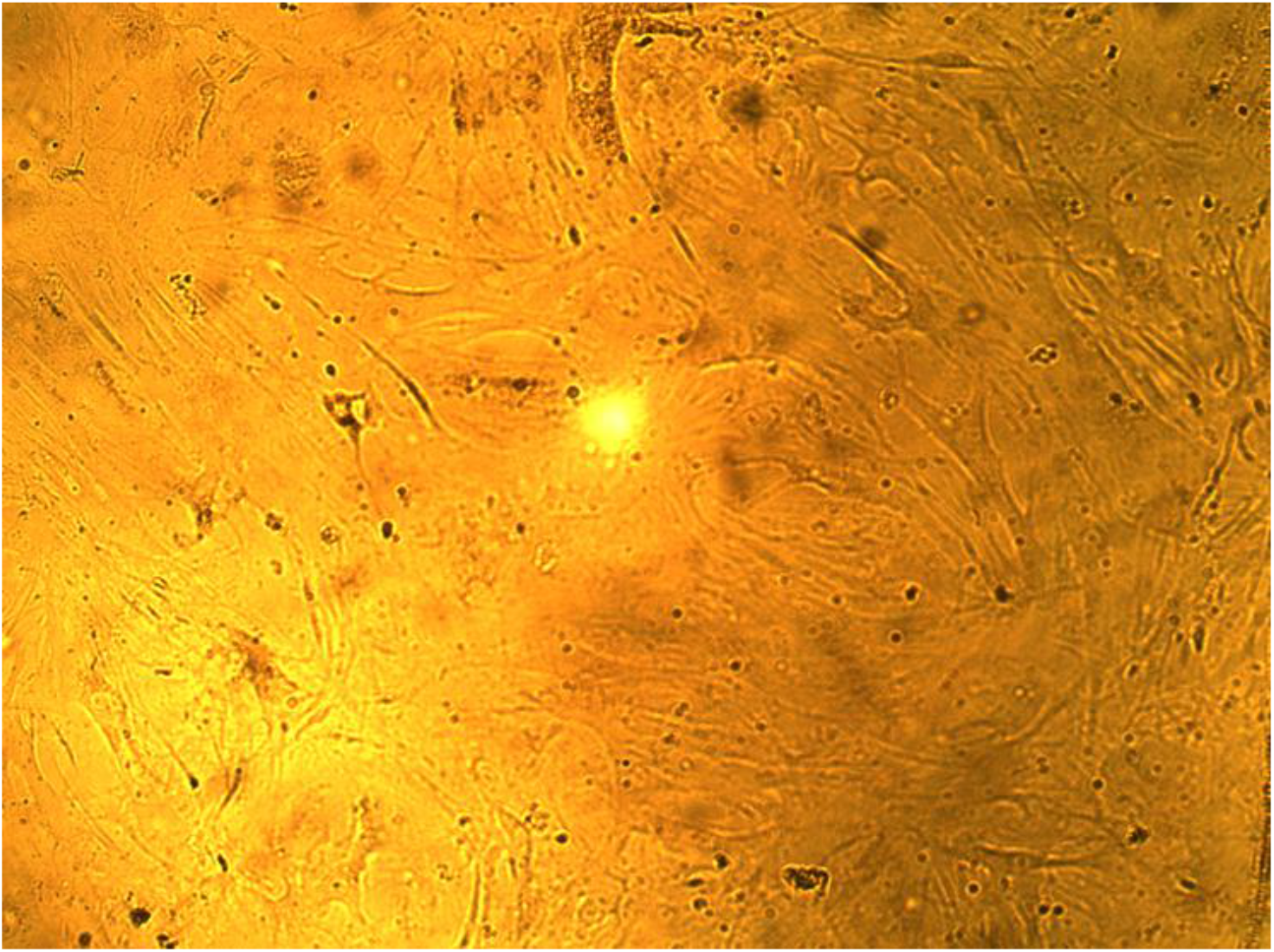
HDSF hTERT 55 passage, growth 40 days after re-seeding.

**Figure 11.**
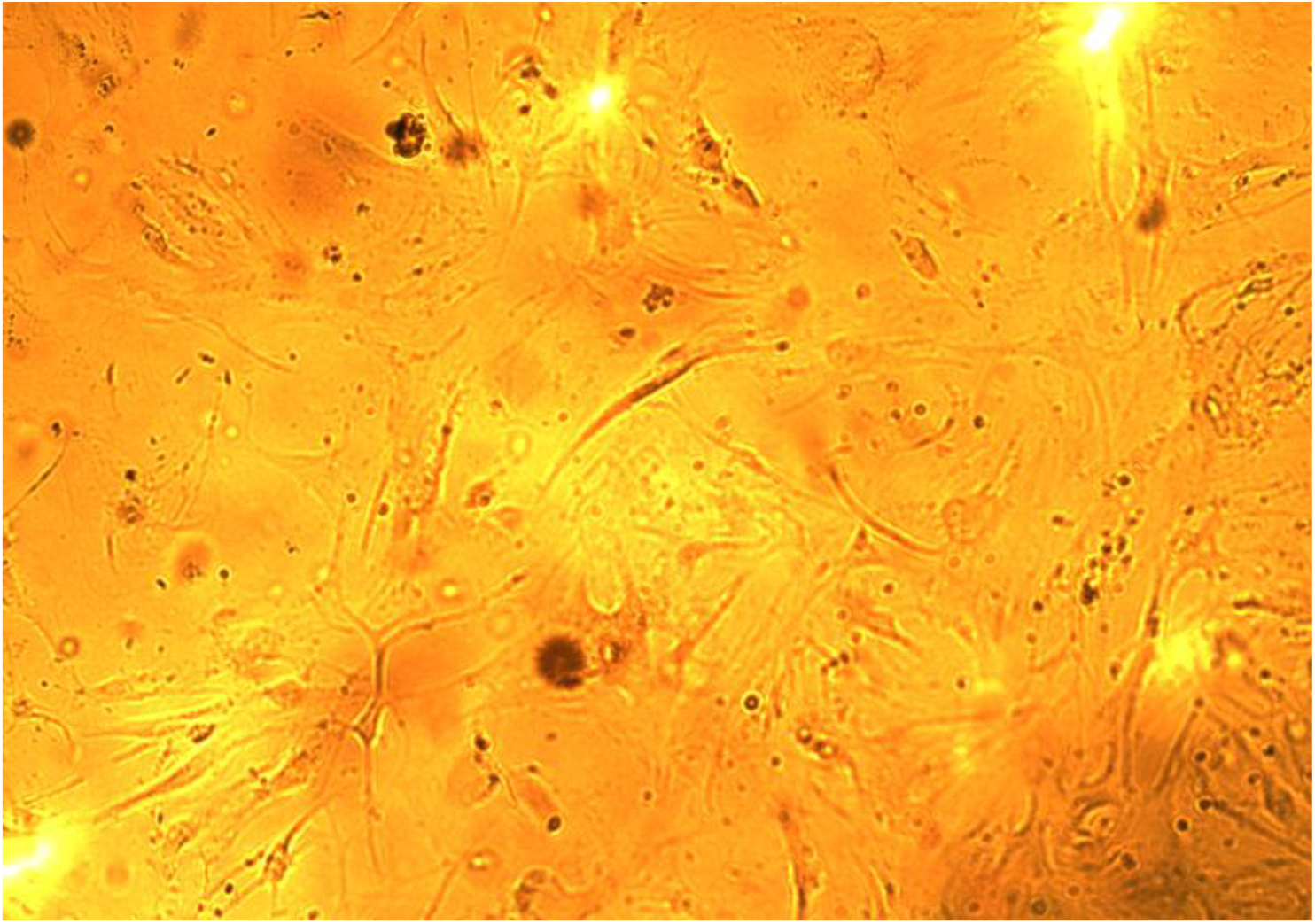
HDSF hTERT 57 passage, growth 33 days after re-seeding.

Telomerized and non-telomerized cells also have differences in appearance at the late passages. Non-thelomerized cells are more compact and more rounded, as can be seen in Figure 12, which shows the cells in the 47th passage after 31 days of growth after re-seeding. Telomerized cells are shown in the previous Figure 11 on the 57th passage after 33 days of growth − they are more elongated and have many outgrowths.

**Figure 12.**
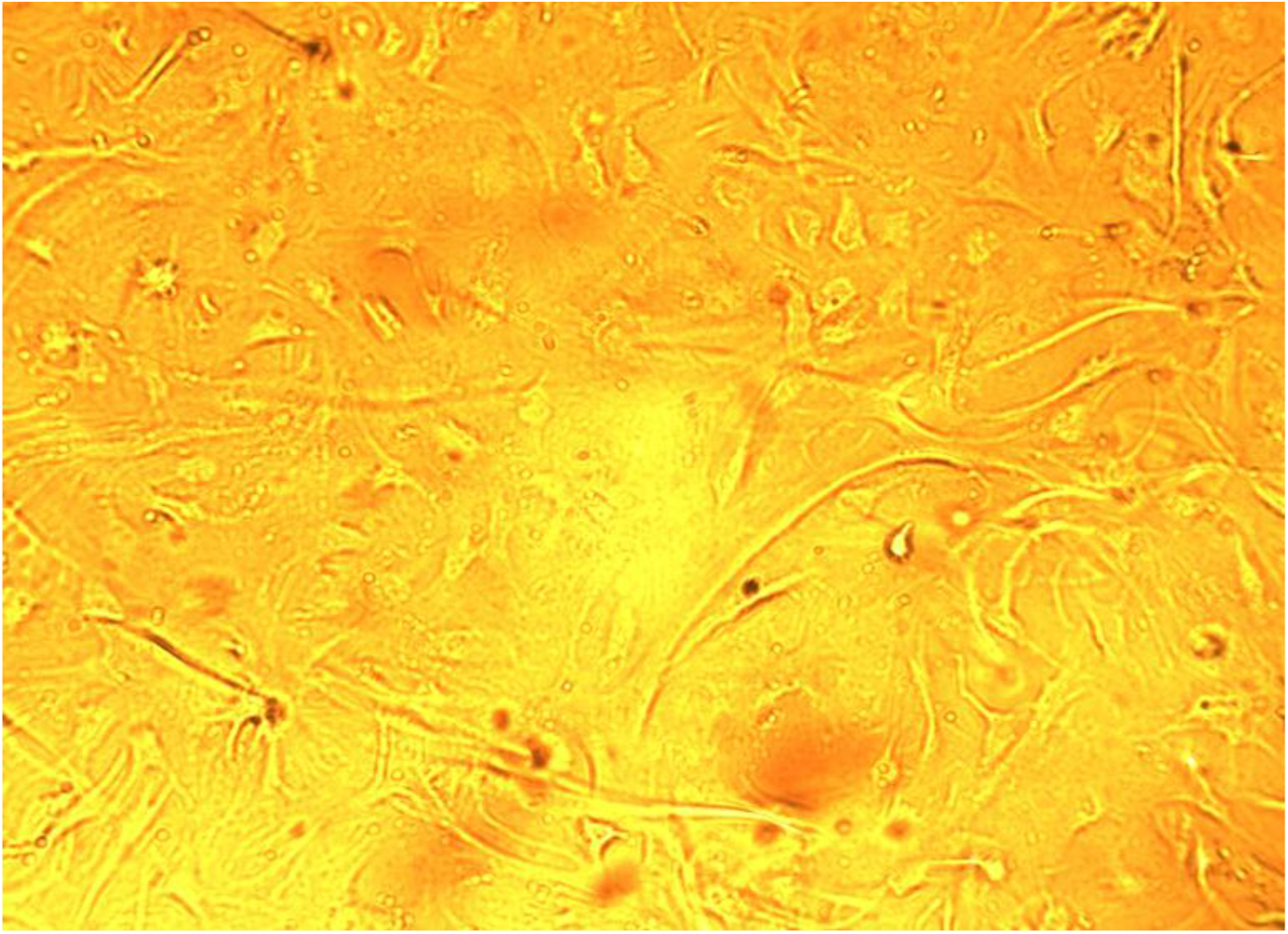
HDSF 47 passage, growth 31 days after re-seeding.

The control cells on the 47th passage and the experimental cells with gene hTERT on the 57th passage practically stopped dividing and the number of cells does not increase. Cultures are observed in this state for more than 245 days. Once every 1-2 weeks, the growth medium was changed to a fresh medium. At the same time, the growth medium in flasks with both types of cells eventually becomes acidified (turns yellow) after the next replacement of the growth medium with a new one, the pH shifts to the acidic side, which indicates the conversion of oxygen to carbon dioxide, as well as the presence of metabolism in the cells. The cells in both cultures are gradually destroyed over time.

## Discussion

The results obtained confirm the data of V. J. Cristofalo et al. (1998) that the number of passages of normal skin cells of an adult donor in culture can be comparable to the number of passages of embryonic cells. It remains unclear why cells in the human body during its growht, starting from the zygote and until the end of growth, already pass on average about 44-45 doubling (Prokhorov, 1999), and cells obtained from an adult organism can pass in culture an additional 45-60 divisions (Terekhov, 1984; Egorov et al., 2003; and the data of our article). It turns out that the potential of cell division from the zygote to the end of growth and plus the number of cell divisions in the culture is already (in the sum) a total of 89-105 doubling. Since cell division in the body almost stops after the end of its growth, it turns out that the cellular potential of somatic cells is not exhausted and is, at least for skin fibroblasts, the same 45-50 doubling. This provides food for thought, why this happens and suggests the possibility of using the unused cellular potential in the body to solve problems with organ health and to fight aging.

## Conclusion

After a certain number of passages, cell division in culture slows down significantly. The cells of the control cultures without the telomerase gene still divide at 46-47 passages, but they are much slower and have a “ bad “ appearance. Therefore, to update the cell mass in cell therapy, it is desirable to use cells with a sufficient rate of division and having a good appearance up to about 25-30, a maximum of 33-35 passages.

At large passages (more than 40), the appearance of the cells changes greatly, they become larger, there are many outgrowthes up to 6-7-10 in number, and the cell area increases. The doubling time of the culture increases after the 27th passage. Telomerized cells pass 10 passages more than control cells. At the same time, the rate of division of telomerized cells persists longer than in control cultures, due to this, telomerized cells overtake control cultures with a time of 9-10 passages.

The rate of cell division in both cultures may additionally depend on the quality of the serum used in the growth medium.

Judging by the results of the experiment, only partial telomerization of adult skin cell cultures took place, because although the cells undergo a greater number of passages compared to the control ones, their division rate still slows down later.

In our experiment, the life span of the control cell culture according to Hayflick (the culture does not double in 2 weeks) is 42 passages, and the telomerized one is 48 passages. In fact, the control culture doubles at least 5 more times after the Hayflick “death”, and the culture with telomerized cells doubles 9 times.

It can be concluded that the division potential of non-telomerized normal cells is sufficient for the restoration of old tissues and organs, and the main obstacle to the implementation of rejuvenation of the adult body is the old cells and structures of tissues and organs of the body. To eliminate old cells and structures, it is necessary to select substances such as enzymes that can do this, and the remaining young cells will be able to divide and renew the tissues and organs of the body. Finding out the limits of cell proliferation of intact somatic cells, as well as cells with an embedded telomerase gene, will allow better use of cell cultures to restore old organs and tissues, counteract aging and increase human life expectancy (Prokhorov, 2021).

Telomerized cells will need to be used for the rejuvenation of old or the treatment of diseased organs in the event that the cells of these organs have exhausted their usual proliferative potential.

This research did not receive any specific grant from funding agencies in the public, commercial, or not-for-profit sectors.

